# Engineering large chromosomal deletions by CRISPR-Cas9

**DOI:** 10.1101/2020.12.29.424675

**Authors:** Thomas F. Eleveld, Chaimaa Bakali, Paul P. Eijk, Phylicia Stathi, Pino J Poddighe, Bauke Ylstra

**Affiliations:** Department of Pathology, Amsterdam University Medical Centers, Cancer Center Amsterdam, De Boelelaan 1117, 1081 HV Amsterdam, the Netherlands; Department of Clinical Genetics, Amsterdam University Medical Centers, Cancer Center Amsterdam, De Boelelaan 1117, 1081 HV Amsterdam, the Netherlands

## Abstract

Arm-level chromosomal deletions are a prevalent and defining feature of cancer. A high degree of tumor-type and subtype specific recurrencies suggest a selective oncogenic advantage. However, due to their large size it has been difficult to pinpoint the oncogenic drivers that confer this advantage. Suitable functional genomics approaches to study the oncogenic driving capacity of arm-level deletions are limited. Here we present an effective technique to engineer arm-level deletions by CRISPR-Cas9 and create isogenic cell line models. We simultaneously induce double-strand breaks (DSBs) at two ends of a chromosomal arm and select the cells that have lost the intermittent region. Using this technique, we induce arm-level deletions on chromosome 11q (65 MB) and chromosome 6q (53 MB) in neuroblastoma cell lines. Such isogenic models enable further research on the role of arm-level deletions in tumor development and growth and their possible therapeutic potential.

## Introduction

Somatic genetic alterations, like mutations, translocations and copy number changes, are the driving force of cancer, involved in not only oncogenesis, but also tumor evolution and progression, therapy sensitivity and resistance^1^. Genetic alterations are tumor type and even subtype specific and determine their malignant potential. Functional genomics is a mainstay to comprehensively study the role of the genetic alterations to explain tumor development and growth.

Gene specific mutations have given a lot of insight into the oncogenic processes driving certain cancer types. For a variety of such mutations these studies have led to clinically effective treatment modalities that directly inhibit the function of the mutant proteins, such as the use of specific BRAF inhibitors in tumors with BRAF mutations^2^. Translocations that recurrently affect particular genes are relatively easily modeled and have therefore been extensively studied. For several translocations this has also led to precision medication that targets the fusion protein, like Imatinib for tumors that harbor the BCR-ABL oncogene^3^. Copy number changes come in a large size range from focal events, affecting small regions of up to 3 MB with 1 or few genes, to entire chromosomes, chromosomal arms or large parts thereof, affecting hundreds to thousands of genes^4^. For recurrent focal chromosomal copy number aberrations the driving genes are also relatively easily identified, exposing strong driving oncogenes and tumor suppressor genes^5^, and also in some cases other functional elements like microRNA’s^6^. Several of these can also be targeted by specific inhibitors, leading to significant clinical benefit, like for the use of Herceptin for HER2 amplified cancers^7^. For most recurrent arm-level chromosomal aberrations, however, the large number of genes and regulatory elements affected complicates the elucidation of the underlying oncogenic processes. Subsequently, no targeted medicine is available yet that target arm-level copy number aberrations.

Arm-level chromosomal copy number aberrations can be highly recurrent, with the most frequent events, like 8p loss, 17p loss and 8q gain, occurring in >30% of solid malignancies^8^. Arm-level events affect a vastly larger number of genes than other genomic changes (mutations, translocation, focal events), but each gene to a much smaller extent. This observation, combined with the paucity of clear driving genes or regulatory elements behind these arm-level changes, has led to the suggestion that arm-levels aberrations could possibly promote tumor growth not by affecting the function of a single driver, but by concerted regulation of multiple genes and regulatory elements located on the involved region^9,10^

The molecular function of most established oncogenes and tumor suppressor genes has been extensively studied through the use of functional genetic experiments in relevant model systems (knockdown, knockout, overexpression etc.). However, very little is known about the functional effects of arm-level chromosomal copy number aberrations, since it is difficult to study the effects of subtle changes in the expression of multiple genes by conventional molecular biology approaches. The few studies that have addressed this subject have shown counterintuitive results, demonstrating that gain of single chromosomes is associated with a tumor-suppressive rather than tumor-promoting effect^11,12^.

Here we present a technique to induce large chromosomal deletions using CRISPR/Cas9 by simultaneously introducing double strand breaks (DSBs) at two locations within one chromosomal arm and a synthetic ssDNA template that spans the created gap for repair. Using this protocol, arm-level deletions that are observed in patients and are associated with poor prognosis are introduced, in neuroblastoma cell lines. Terminal deletions on chromosome 11q (65MB) and 6q (53 MB), were modeled by making breaks and inducing translocations between sites where deletions were recurrently observed in patients and the telomere of the targeted chromosome, with up to 30% of clones containing the desired deletion. With this technique we offer a method for the generation of isogenic model systems of arm-level chromosomal deletions.

## Results

### Engineering an 11q deletion in neuroblastoma SKNSH cells

Terminal deletions affecting a large part of chromosome 11q are frequently observed in neuroblastoma patients and are associated with a very poor prognosis^13^. The high degree of recurrence (34%) and link to poor prognosis suggest an oncogenic effect, however the underlying molecular pathways that drive this effect remain to be elucidated. Therefore, we aimed to establish an isogenic model of such a deletion in the neuroblastoma cell line SKNSH, which is diploid with two intact copies of chromosome 11^14^. We hypothesized that DSBs at two locations, one at chromosome band 11q13.4 and the other at chromosome band 11q25 near the telomere, could lead to joining of the two ends with deletion of the intermediate region. Guide RNAs were designed to induce DSBs within 100bp of the 11q breakpoint from the neuroblastoma cell line GIMEN, which has lost the terminal regions of 11q through a der(11)t(11;17)^15^, and near the telomere on 11q (Figure 1A,B). Joining of these DSBs would induce the same loss of 11q in the cell line SKNSH as observed in the GIMEN cell line, apart from the 11q telomeric region that would remain in SKNSH.

**Figure 1.**
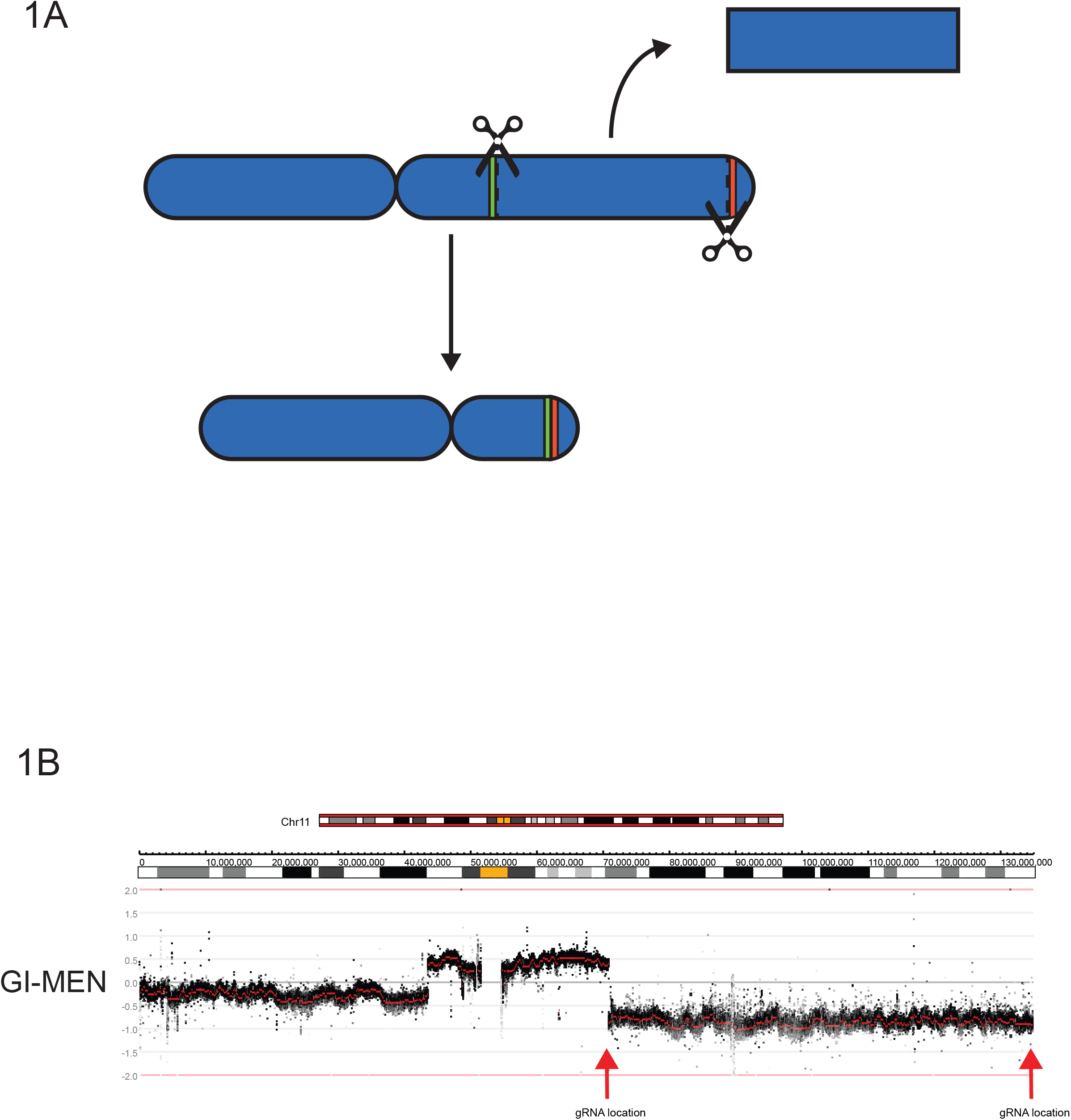
The concept of CRISPR-induced chromosomal deletions. **A)** Schematic overview of chromosome 11. Scissors represent the gRNA locations and green and red colors the regions adjacent to the gRNAs. DSBs could lead to joining of the green and red region with loss of the intermittent region. **B)** Copy number profile of chromosome 11q of the neuroblastoma cell line GIMEN. Data was generated by WGS and visualized using the R2 bioinformatics platform. Black dots represent values for bins and red dots represent segmented values. The red arrows represent the gRNA location at chromosome 11q13.4and near the telomere at 11q25. Ideograms representing the chromosomal location and banding pattern (centromeres are represented in yellow) are shown above the profile.

Complexes consisting of Cas9-protein and synthetic gRNA’s (Ribo Nucleo Protein, RNP complexes) were introduced into cells using electroporation. Single RNP complexes have a 60-70% editing efficiency as determined by Inference of CRISPR Edits (ICE) analysis (Synthego Performance Analysis, ICE Analysis. 2019. v2.0. Synthego, Supplementary Figure 1). To facilitate joining of the two DSBs a 100 bp single stranded DNA (ssDNA) molecule consisting of 50 bp stretches of homology to the regions surrounding the DSBs was added to the reaction. To determine whether the desired end-joining took place, PCR was performed on DNA isolated from the bulk transfected cells using primers spanning the expected translocation regions (Figure 2A, Supplementary Figure 2). Gel electrophoresis of the PCR reaction shows bands at the expected size, indicating the presence of a fusion product, while Sanger sequencing confirmed the specificity of the PCR product. Control primers located in close chromosomal proximity confirmed the integrity of the DNA and showed that one wild-type copy of chromosome 11 was still present. Interestingly, the translocation PCR product was not observed when the homologous ssDNA was omitted, indicating that it aids in joining the two DSBs with loss of the intermittent region.

**Figure 2.**
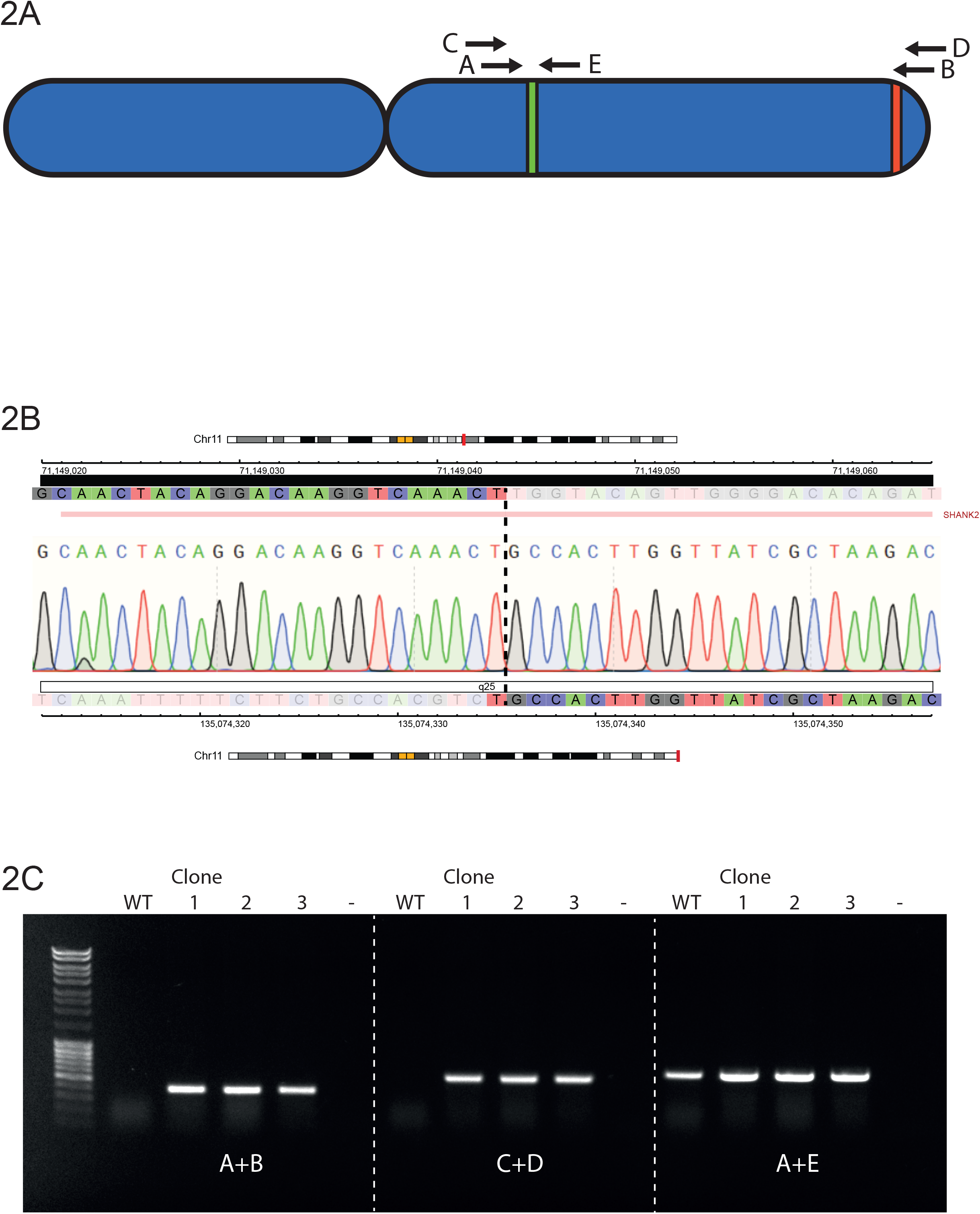
PCR analysis indicates the presence of a translocation between 11q13.4 and 11q.25 **A)** Schematic overview of chromosome 11, with green and red regions representing the regions adjacent to the gRNA recognition sites and letters representing primers used for analysis. **B)** Sanger sequencing electropherogram results of the translocation product from SKNSH 11q clone 4. Color coded tracks above and below the trace represent the sequence (C=blue, T=red, G=grey, A=green) and chromosomal location of the first and second part, respectively. Ideograms representing the chromosomal location, banding pattern (centromeres are represented in yellow) and genes located in the two regions are shown above and below the profile **C)** Agarose gel electrophoresis of the PCR reaction of the wild-type (WT) and the three clones (1, 2 and 3) from two experiments that were positive for the presence of a translocation. The first lane contains Generuler 1kb DNA ladder (Thermo Fisher Scientific). Primers combinations are shown below the bands and refer to the primers shown in 2A.

To obtain a homogenous population with the engineered chromosomal loss, after four passages cells were seeded at low density (2000 cells per 10 cm Ø plate) to derive single clones. DNA was subsequently isolated from these clones and the presence of a translocation product was screened using the previously described PCR strategy. The four first clones that reached a cell density that allowed us to isolate sufficient DNA, all contained a translocation PCR product and Sanger sequencing showed the same sequence for all clones (Figure 2B). However, the sequenced PCR products did not completely match the ssDNA template sequence which would have been expected if the template had been instrumental in the DSB repair. All four clones contained the same 15 bp deletion compared to the anticipated template sequence. Since these cells were cultured for a few passages after electroporation, we concluded that these clones are most likely derived from a common ancestor and are not independent. To determine the reproducibility of the procedure and establish the frequency at which these translocations occur, the 11q deletion engineering experiment was performed again with a new passage of the SKNSH cell line. However, to prevent early outgrowth of positive clones due to possible selective advantages, cells were seeded at lower, density (1000 cells per 15 cm Ø plate) 72 hours post-electroporation. PCR products spanning the intended fusion were detected in two out of nine clones (Supp Figure 3). One of the two positive clones (clone 7) showed a ~350 bp duplication of the 11q13.4 region adjacent to the gRNA recognition site, while the other positive clone (clone 10) showed a 23bp insertion (data not shown), which shows they are of independent origin. The PCR verification of the two independent positive clones from this experiment and the positive clone from the previous experiment are shown in Figure 2C.

**Figure 3.**
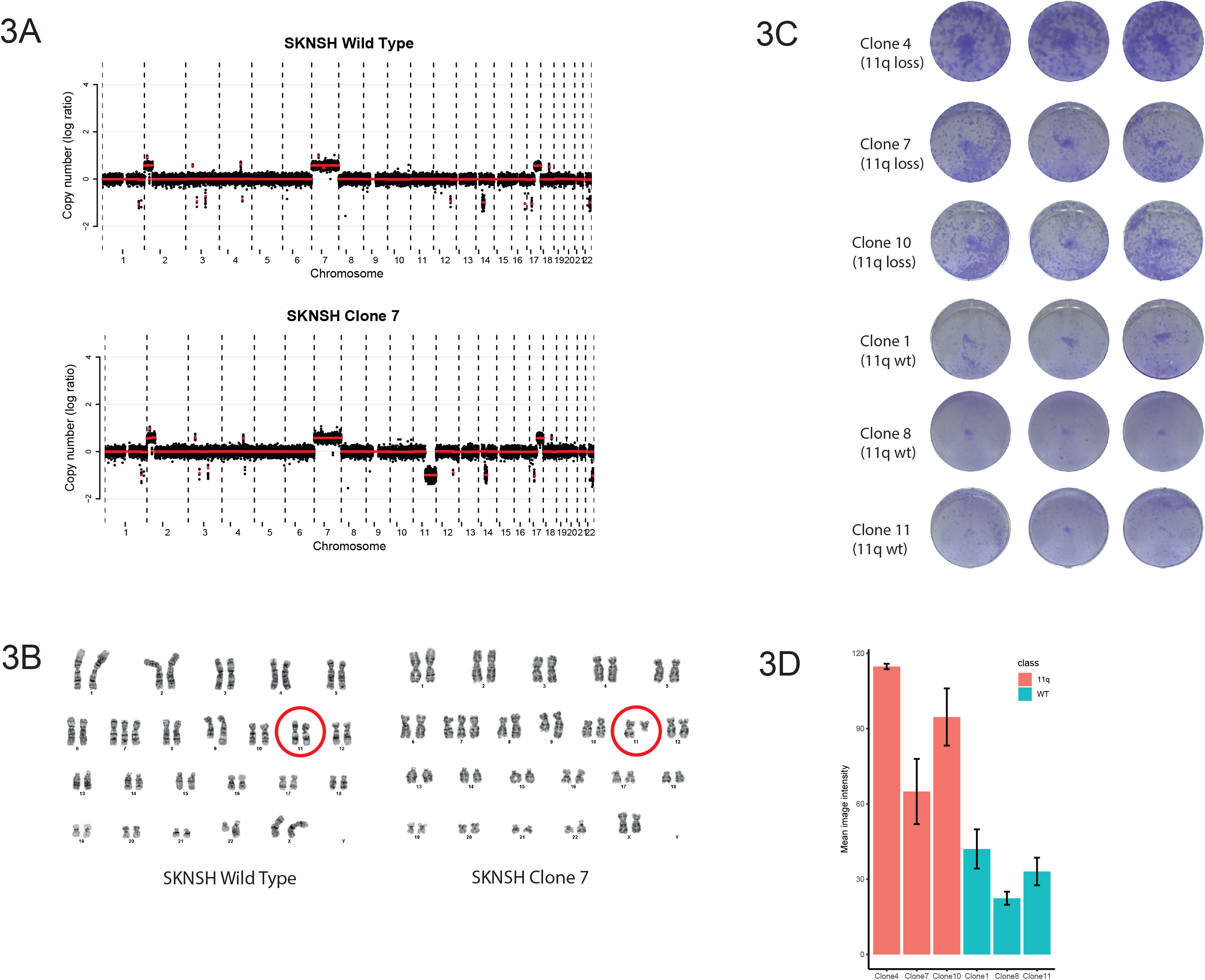
CRISPR induced 11q deletions lead to increased clonogenic capacity in SKNSH **A)** Copy number profiles of wild type SKNSH cells and chromosome 11q deletion clone 7 (one of the clones that was positive for the translocation PCR product). The x-axis shows the genomic location and the y-axis shows median-normalized log2-transformed copy number, with black dots representing bins and red lines representing segmented copy numbers. **B)** Karyotypes of the wild type cells and chromosome 11q deletion clone 7 cells. Red circles indicate the chromosome 11 pair. Karyotype of the wild type cell line: 47,X,add(X)(p21.2),+7,add(9)(q34.1),add(22)(q13). Karyotype of the clone: 47,X,add(X)(p21.2),+7,?add(8)(p23),add(9)(q34.1),del(11)(q13),add(22)(q13) **C)** Crystal violet staining of colony forming assays of three clones with 11q deletion and three clones without. **D)** Quantification of the experiment shown in 3D. Error bars represent the standard deviation of the three wells for each line. Pairwise t-test analysis shows significant differences between all chromosome 11q deletion clones and all negative clones.

The three clones with a translocation PCR product were analyzed for genome-wide copy number aberrations using shallow Whole Genome Sequencing (shallow WGS), together with the wild-type cell line and the three clones that that did not show a translocation PCR product. All three clones with a translocation product showed a clear loss of one copy of the intended region of chromosome 11q, while the other three clones and the wild-type did not show this arm-level loss (Figure 3A, Supplementary Figure 4). Across the entire genome, no additional chromosomal copy number differences were observed for the three PCR positive clones, compared to the PCR negative clones or the parental cell line. Shallow WGS offers a high resolution to detect copy number aberrations, yet translocations cannot be detected. To determine whether no additional, copy number neutral, translocations occurred due to the CRISPR editing, classical karyotyping was performed for the wild-type parental cell line and PCR positive clone 7. A terminal deletion of chromosome 11 with breakpoint at 11q13 was the only karyotypical difference between this clone and the wild-type (Figure 3B). We conclude that the CRISPR-Cas9 genome editing has indeed caused a translocation between the two regions where DSBs were induced with concomitant deletion of the interstitial region. This cell line, together with the wild-type, represent an isogenic cell line pair that can serve as a model to study the effects of 11q loss in neuroblastoma.

**Figure 4.**
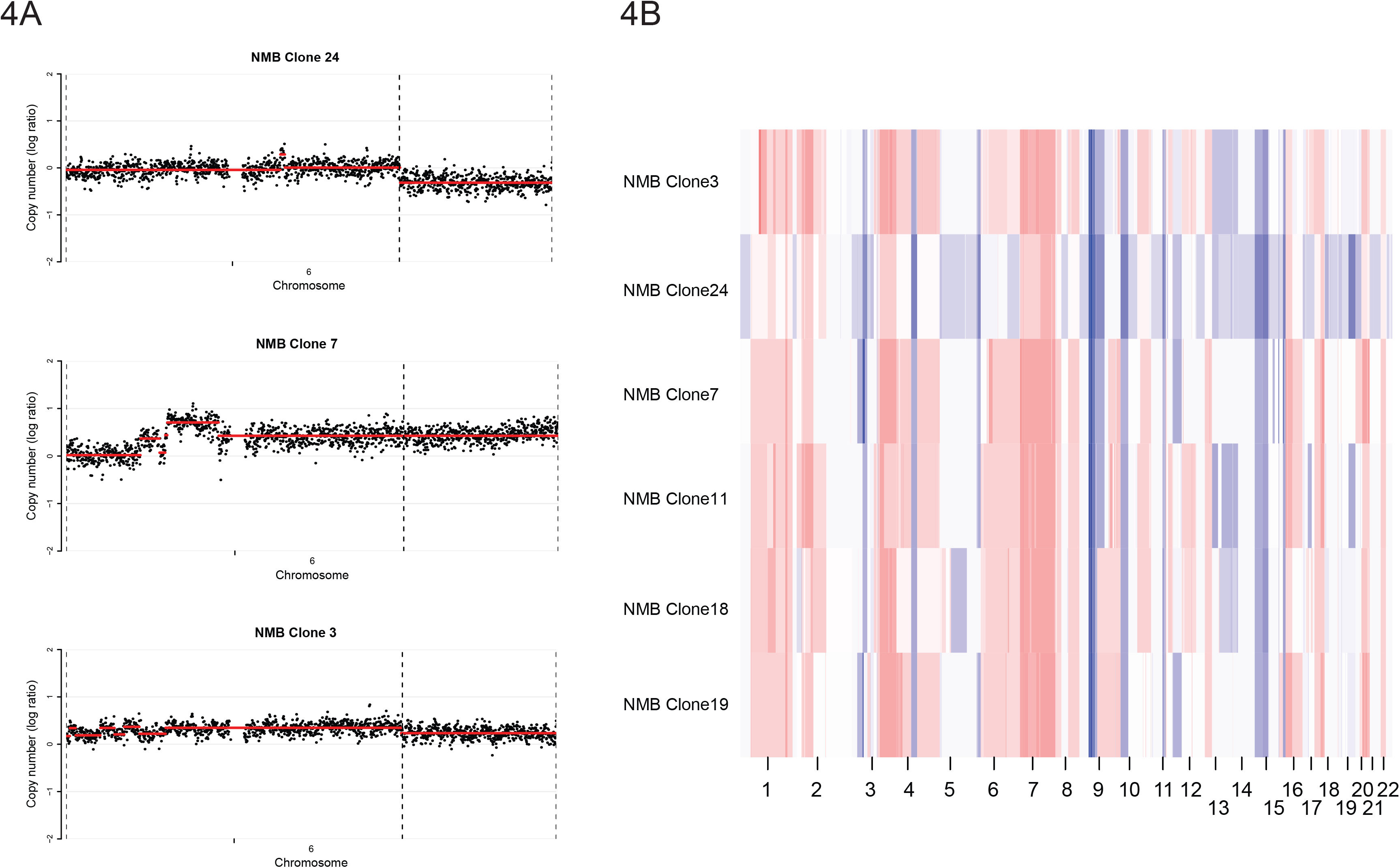
Chromosome 6q deletion and genomic heterogeneity in the cell line NMB **A)** Copy number profiles of chromosome 6 of the three clones that were positive for a translocation PCR product. The x-axis shows the genomic location, and the y-axis shows median-normalized log2-transformed copy number, with black dots representing bins and red lines representing segmented copy numbers. Clone 24 and 3 show a deletion on 6q, while this is not observed in clone 7. **B)** Whole genome overview of the copy numbers of all NMB clones. Each horizontal bar represents the copy number profile of one clone on each chromosome (shown on the x-axis). Colors represent the copy numbers in each region, ranging from blue (log2 ratio < 0), to white (log2 ratio = 0), to red (log2 ratio <0).

SKNSH clones with chromosome 11q deletions were observed at a high frequency (2/9, 22%) without the use of selection. This led us to hypothesize that chromosome 11q deletion, observed in 34% of neuroblastoma tumors^13^, might give a selective advantage. The SKNSH clones with the translocation were among the first colonies to reach sufficient cells to be harvested for DNA isolation in both experiments (data not shown), suggesting that loss of the long arm of chromosome 11 might cause an increased colony forming capacity. To determine whether this is the case, the same six clones that were characterized by shallow WGS were plated at low density and were stained with crystal violet after 14 days. Clones with 11q deletions showed significantly higher colony forming capacity, as determined by the mean image intensity of the stained wells, than clones that underwent the same procedure but did not show a deletion (Figure 3C, 3D), suggesting that 11q loss indeed provides an advantage in clonal outgrowth.

To determine whether this experimental setup is also effective for other chromosomal deletions in other cell lines we aimed to introduce a 6q deletion in the neuroblastoma cell line NMB. Distal 6q loss is observed in 5.9% of neuroblastoma patients and is associated with extremely poor prognosis^16^. Similar to the 11q experiments, guide RNAs were designed at two locations on chromosome 6q; one at 6q22.1 in close proximity to a relapse specific 6q deletion^15^ (Supplementary Figure 5) and the other at 6q27 close to the telomere. Since the ssDNA used in the 11q experiment was not used for repair, as determined by Sanger sequencing, we omitted a ssDNA template from the experimental setup for chromosome 6q. Cells were transfected with the RNP and DNA was isolated from part of the transfected cells. A translocation PCR product was already observed in, these bulk transfected cells (Supplementary Figure 6), and single cell clone selection yielded three out of the nine clones positive for such a product (Supplementary Figure 7). Shallow WGS showed a deletion in two of these clones, while one clone with a translocation PCR product and three clones without a translocation product did not show loss of this region (Figure 4A, Supplementary Figure 8). Three clones that did not show a translocation PCR product did not show loss of chromosome 6q. In general, it becomes clear that the clones, both with- and without a 6q deletion, have a very different chromosomal make-up. They show major differences in copy numbers of chromosomes other than 6q and even apparent ploidy changes (Figure 4B). Whether this genomic instability is the result of the CRISPR-Cas9 editing or is inherent to the cell line used remains to be determined. Notwithstanding, the described CRISPR procedure allows for a fast and efficient targeted deletion, also for chromosome 6q.

In conclusion, we have successfully developed a procedure to efficiently induce large chromosomal deletions in neuroblastoma cell lines using CRISPR-Cas9. It is reasonable to assume that this method will be applicable to other cancer types and other chromosomes which would enable the rapid generation of isogenic cell models that would allow the study of molecular mechanisms that drive clinically relevant chromosomal deletions.

## Discussion

Recurrent arm-level chromosomal copy number aberrations are observed in almost all cancer types and are often associated with poor prognosis^17,18^. Despite the observation that the most recurrent events occur at similar frequencies as the most frequently mutated gene, TP53^19^, and more frequently than the most frequent focal copy number aberrations^1^, their molecular effects have been far less studied and consequently less understood. Therefore, we devised a strategy to induce arm-level chromosomal deletions using CRISPR-Cas9 by introducing DSBs at known breakpoints and telomeric regions and providing a ssDNA template for repair. The presented method is easy and fast, and we observed high frequencies with more than 30% of clones with the intended chromosomal deletion.

Recently, three alternative methods have been described that also enable the modelling of large chromosomal rearrangements. CRISPR-Cas9 has already been employed to fuse two DSBs on two different chromosomes together for the generation of a fusion gene. However, here a selectable marker was used to select cells with the designed fusion and no copy number aberrations were involved^20^. In another study, TALENs were used to engineer a chromosome 8p deletion in breast cancer cells^21^. This approach was very similar to ours, apart from the use of TALENs instead of CRISPR-Cas9 technology. Finally, CRISPR-Cas9 was used to delete chromosome 3p in lung cells, however, here DSBs were induced near the centromere, but combined with an artificial telomere plasmid with a selectable marker^8^. The advantage of our method is that no molecular cloning is involved, since all reagents can be designed *in silico* and readily ordered, making it cheaper and faster than the alternative methods. Moreover, no exogenous DNA sequences are introduced into the genome of these cells, which makes it a more isogenic model system.

We observe clones positive for the intended chromosomal deletions at high frequency (over 30%). although these frequencies may be imprecise due to the low amount of analyzed clones. Even though the editing efficiencies of the single gRNAs observed were as high as 60%, we would regard these frequencies as improbable without a positive selective effect of the induced deletions on the growth and selection of these colonies for PCR evaluation. In line with this, for chromosome 11q loss in the neuroblastoma cell line SKNSH we observe an increase in colony forming capacity in the 11q deletion clones compared to clones without the 11q deletion. This also explains why the first colonies that had grown to sufficient cell numbers for PCR evaluation contained the intended deletions. This positive effect on the growth of this cell line is contrary to earlier results where chromosomal deletions were induced ^8,21^, where no or little positive effect on proliferation was observed. One key difference is, however, that we use established cancer cells while in these studies karyotypically normal cells were used. It is conceivable that cancer cells already fulfill some of the requirements, i.e. reduced genome maintenance or other hallmarks of cancer^17^, that allow the positive effect of these deletions to manifest itself. If no strong growth advantage of the projected deletion is expected a selectable marker could be added to the ssDNA or plasmid DNA with homology arms and a selection marker could be used as a repair template.

In our initial experimental set-up for chromosome 11q, we used a ssDNA template containing homology regions of both locations where DSBs were induced, since it has been suggested that such templates aid in driving repair through homologous recombination^22^. We did observe PCR products suggestive of successful translocations in the cells where a ssDNA template was added, while these, were not observed in cells without this template. However, Sanger sequencing analysis of single cells clones that showed such a product (and later showed to have acquired the 11q loss), yielded sequences that showed deletions or duplications compared to the ssDNA sequence. It is thought that ssDNA aids in the repair of DSBs through homologous recombination, which is generally considered to be an error-free process^23^, suggesting that the ssDNA templates were not used for repair. Therefore, the ssDNA templates were omitted in experiments where 6q loss was induced. We also observed relatively high frequencies of the intended translocation in these experiments, suggesting that ssDNA templates are not necessary to join the two DSBs. However, it remains to be determined whether they can increase the frequency of the correct translocations in this experimental set-up.

Although CRISPR-Cas9 can be used to induce DSBs at specific locations, it has been known to also produce unintended effects on chromosomal make-up^24,25^. In the 11q CRISPR clones of the cell line SKNSH we do not observe additional large chromosomal changes, apart from the intended 11q deletion. There could be small structural changes or single nucleotide variants that we were unable to detect, however, it seems unlikely that these play a role in the observed increase in clonogenic capacity, since independently derived clones showed the same phenotype. For the cell line NMB we see a very different pattern, with a lot of chromosomal changes besides the intended 6q. It has been established that CRISPR-Cas9 can also cause unintended chromosomal rearrangements and overall changes in ploidy and chromosome number^25^. However, we regard it as unlikely that the large changes that are observed are due to CRISPR-Cas9 genome editing, especially since we did not observe any such changes using the same procedure in SKNSH. It seems more plausible to assume that these results reflect the genetic heterogeneity of the NMB cell line, which may be amplified by the clonal selection step in the procedure.

In conclusion, we show that it is feasible to engineer arm-level chromosomal deletions in a fast and efficient fashion while maintaining the original telomere and without the introduction of exogenous DNA sequences. For the deletion of 11q in the neuroblastoma cell line SKNSH an effect on colony formation was readily observed, which could suggest activation of specific oncogenic pathways. Many studies have addressed potential target genes that confer the oncogenic effect of 11q loss in neuroblastoma^26^, however, this approach has largely failed to elucidate the underlying mechanism or to identify treatment options. Creating isogenic cell lines for chromosomal deletions will obviate this gene-by-gene approach and enable researchers to directly study molecular effects and possible associated therapies with a chromosomal rather than just gene-oriented approach.

## Materials and methods

### Cell culture

Cell lines were cultured in DMEM supplemented with 10% FCS, 20 mM L-glutamine, 10 U/ml penicillin and 10 μg/ml streptomycin and maintained at 37 °C under 5% CO2. Cell line identities were regularly confirmed by short tandem repeat (STR) profiling using the PowerPlex16 system and GeneMapper software (Promega)^27^. Cell lines were regularly screened for mycoplasma.

### Colony formation assay

Single cell suspensions of deletion-positive and deletion-negative clones were seeded into 6-well plates (2 ×10 cells/well). At the endpoints of colony formation assays, cells were fixed, stained with crystal violet and scanned. All clones of each cell line were fixed at the same time. Scans of all wells were quantified using ImageJ software^28^.

### RNP complexes and transfection

Custom crRNA’s, tracrRNA and recombinant Cas9 protein were acquired from Integrated DNA Technologies. The sequences for the crRNA’s are shown in Table 1A. Custom crRNA’s and tracrRNA’s were combined according to the manufacturers’ protocol. RNP complexes were made according to the manufacturer’s protocol in a 6:1 ratio (gRNA: Cas9 protein). Electroporation was performed on 4×10^5^ cells using the Neon transfection system (Thermo Fisher Scientific) with the protocol for SH-Y5Y cells (1200V-20ms pulse – 3 times).

**Table 1A.**
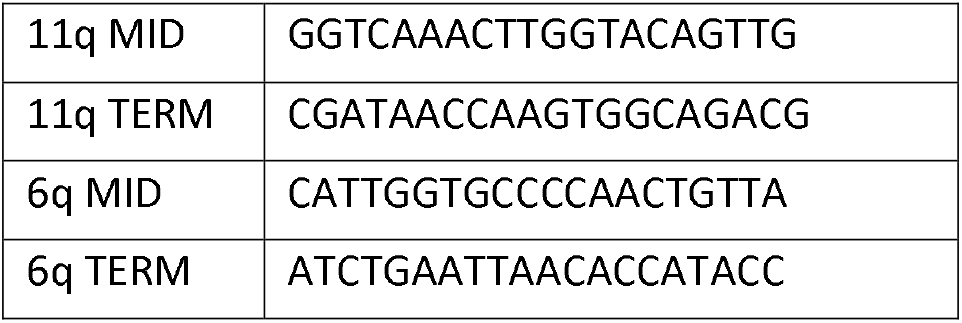
gRNAs used in this study

### Clone selection

Transfected cells were seeded with 1000-2500 cells in a 10/15 cm Ø plate tissue culture plate to form colonies. When colonies reached a size of approximately 5 mm medium was aspirated and colonies were covered with 8 mm cloning cylinders (Dow Corning) coated in sterile 976V silicone high vacuum grease (Dow Corning) to seal it to the plate. Colonies were subsequently trypsinized and transferred to 96 well plates and grown for further characterization and experiments.

### DNA isolation and PCR characterization

DNA was isolated using QuickExtract DNA extract solution (Lucigen) for PCR grade DNA or the QIAamp DNA mini kit (Qiagen) for WGS grade DNA according to the manufacturers’ protocols. PCR was performed using Platinum™ II Hot-Start Green PCR Master Mix (Thermo Fisher Scientific). Primers are specified In Table 1B. Sanger was performed by Macrogen Europe with the same primers used for amplification.

**Table 1B.**
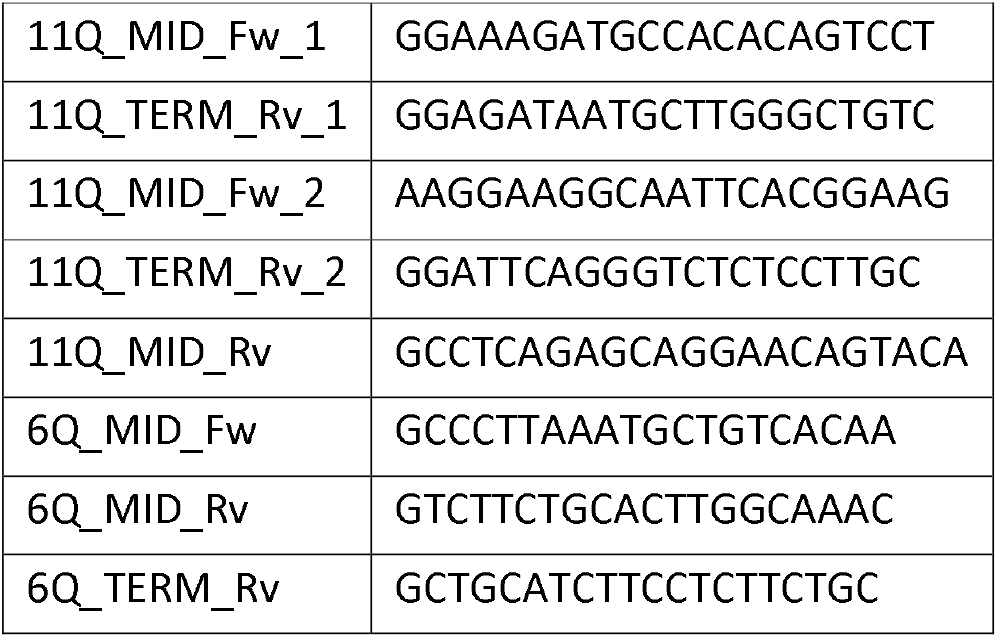
Primers used in this study

### Shallow WGS and bioinformatic analysis

Shallow WGS was performed as described previously^29^. In short, library preparation was carried out, comprising fragmentation of DNA by sonication, repair of DNA ends, 3 prime adenylation, adapter ligation, purification, PCR amplification of the fragments with adapters and lastly again purification. The Illumina Next Generation Sequencing platform was used for shallow WGS. The sequencing data was analyzed using the QDNAseq R-package (v1.26.0) with default settings and 100kbp bins^30^. Correction for sequence mappability and GC content was carried out, as well as filtering of problematic genomic regions. The R2 bioinformatics platform (R2.amc.nl) and R-Studio (v4.0.2) were used for analysis and visualization.

### Karyotyping

Cells were transferred onto Lab-Tek chamber slides (Thermo Fisher Scientific) and grown for 24 hours, the last two hours in the presence of colcemid (Thermo Fisher Scientific). After standard cytogenetic harvesting and Giemsa-Trypsin-Giemsa banding 20 metaphase cells were analyzed from the stimulated culture. The karyotypes were described according to ISCN 2016^31^.

## Notes

### Competing Interest Statement

The authors have declared no competing interest.

