## Supplementary Figures for "Engineering large chromosomal deletions by CRISPR-Cas9"

### Supplementary Figure 1

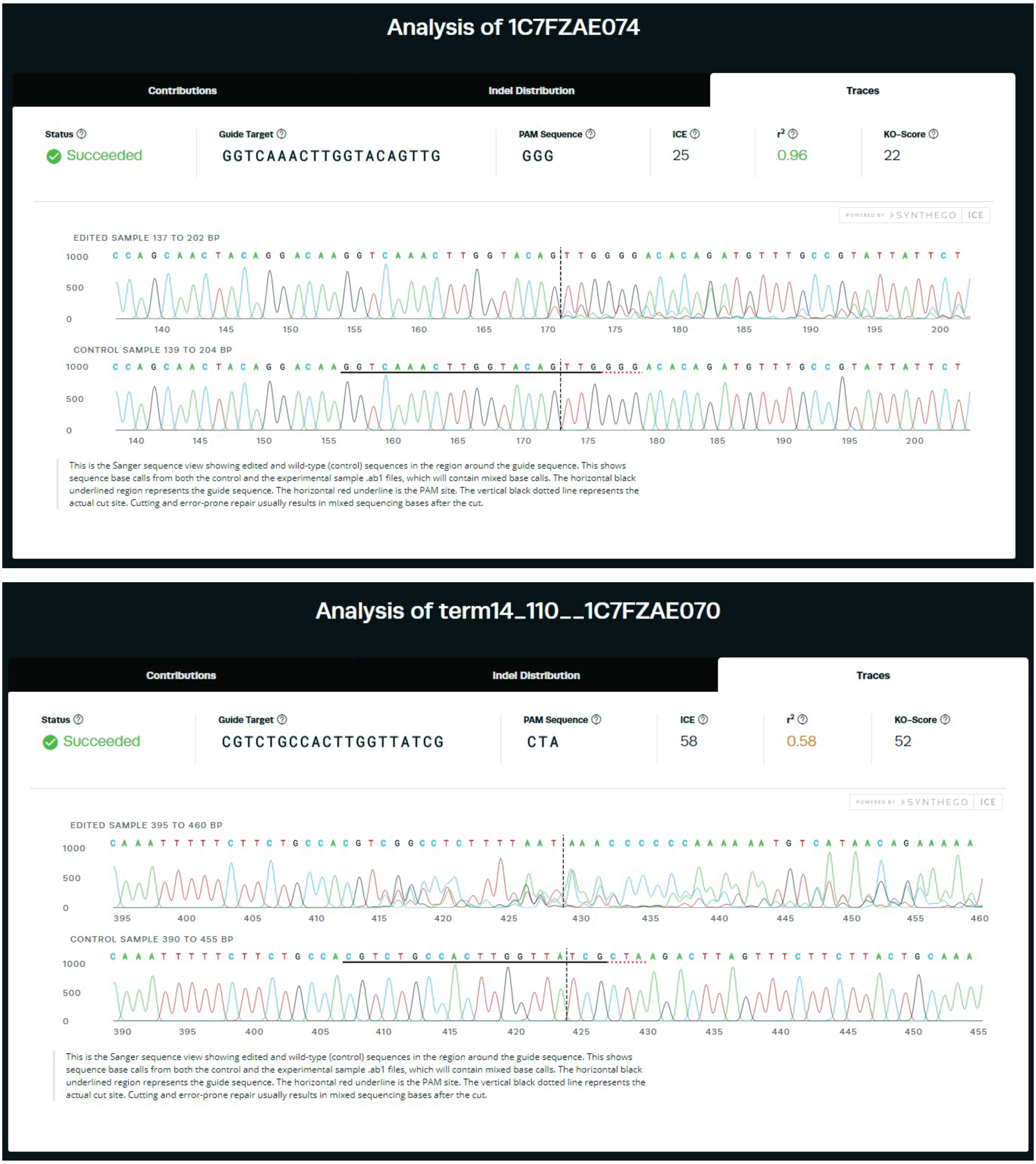


**Supplementary Figure 1:** ICE analysis of efficiency of the 11q MID (upper) and 11q TERM (lower) single gRNAs. ICE scores in the upper bar represent the frequency of insertions/deletions/changes in the edited samples.

### Supplementary Figure 2

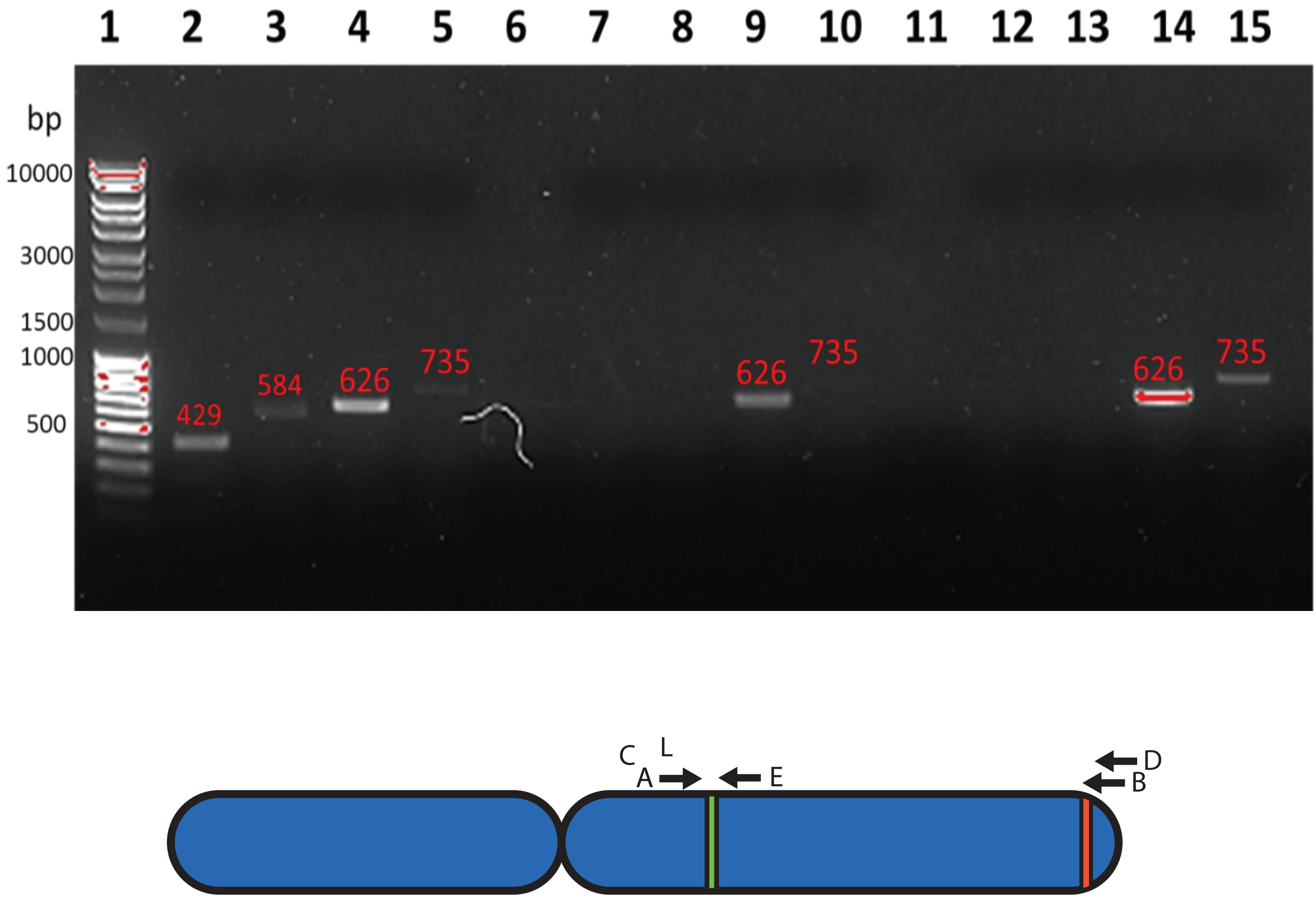


**Supplementary Figure 2:** Agarose gel electropohoresis of PCR reactions performed on cells transfected with 2 gRNAs and a ssDNA (lane 2-5), transfected with 2 gRNAs (lane 7-10) and wild type cells (lane 12-15). Primers combinations are A+B (lane 2,7,12), C+D (lane 3,8,13), A+E (lane 4,9,14) and C+E (lane 5,10,15). Lane 1 contains Generuler 1kb DNA ladder (Thermo Fisher Scientific). Lower figure is duplicated from main Figure 2A for reference.

### Supplementary Figure 3

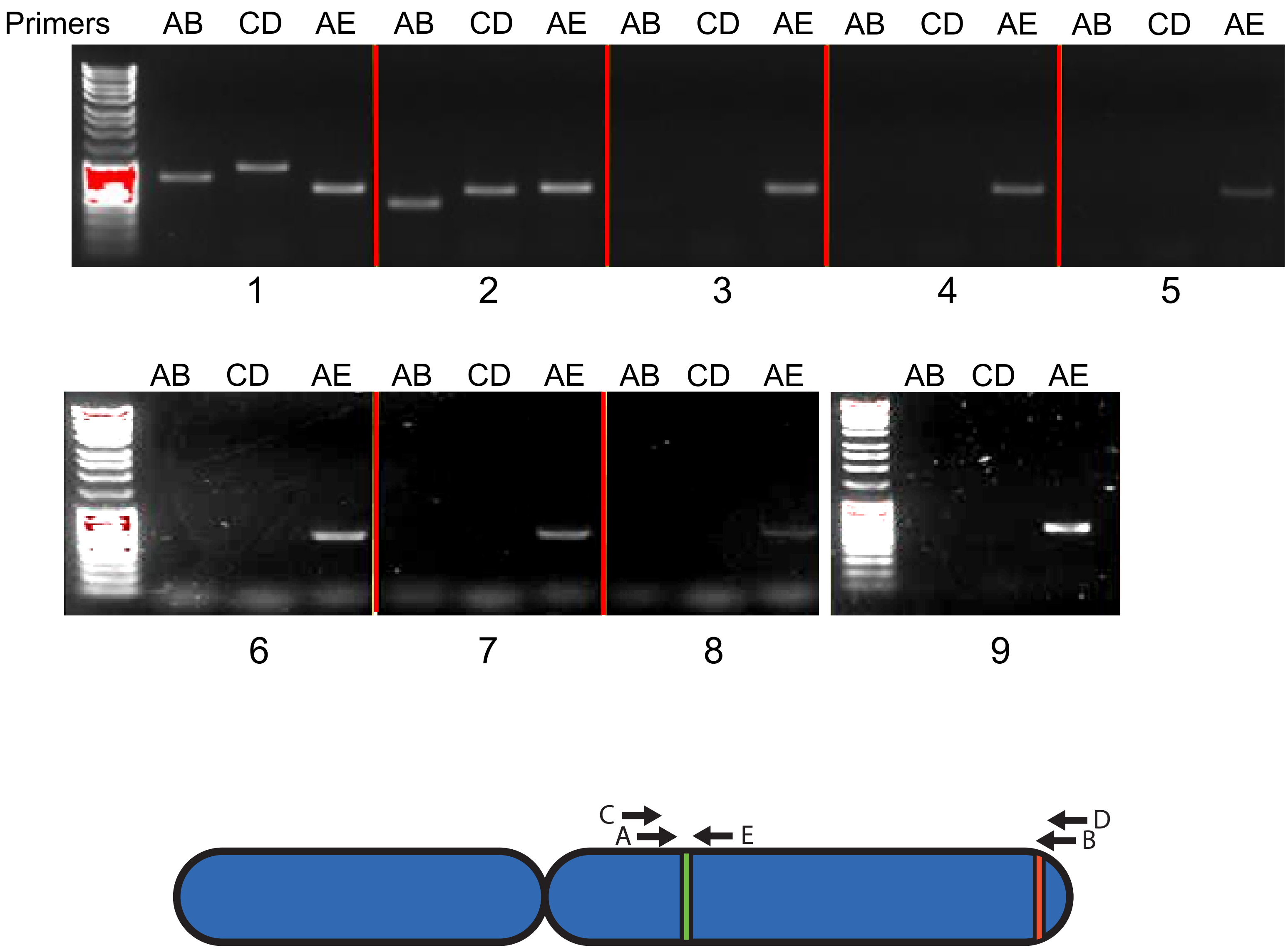


**Supplementary Figure 3:** Agarose gel electropohoresis of PCR reactions performed on 9 clones derived from cells transfected with 2 gRNAs and a ssDNA. Letters indicate the primer combinations used. The first lane of each panel contains Generuler 1kb DNA ladder (Thermo Fisher Scientific). Lower figure is duplicated from main Figure 2A for reference.

### Supplementary Figure 4

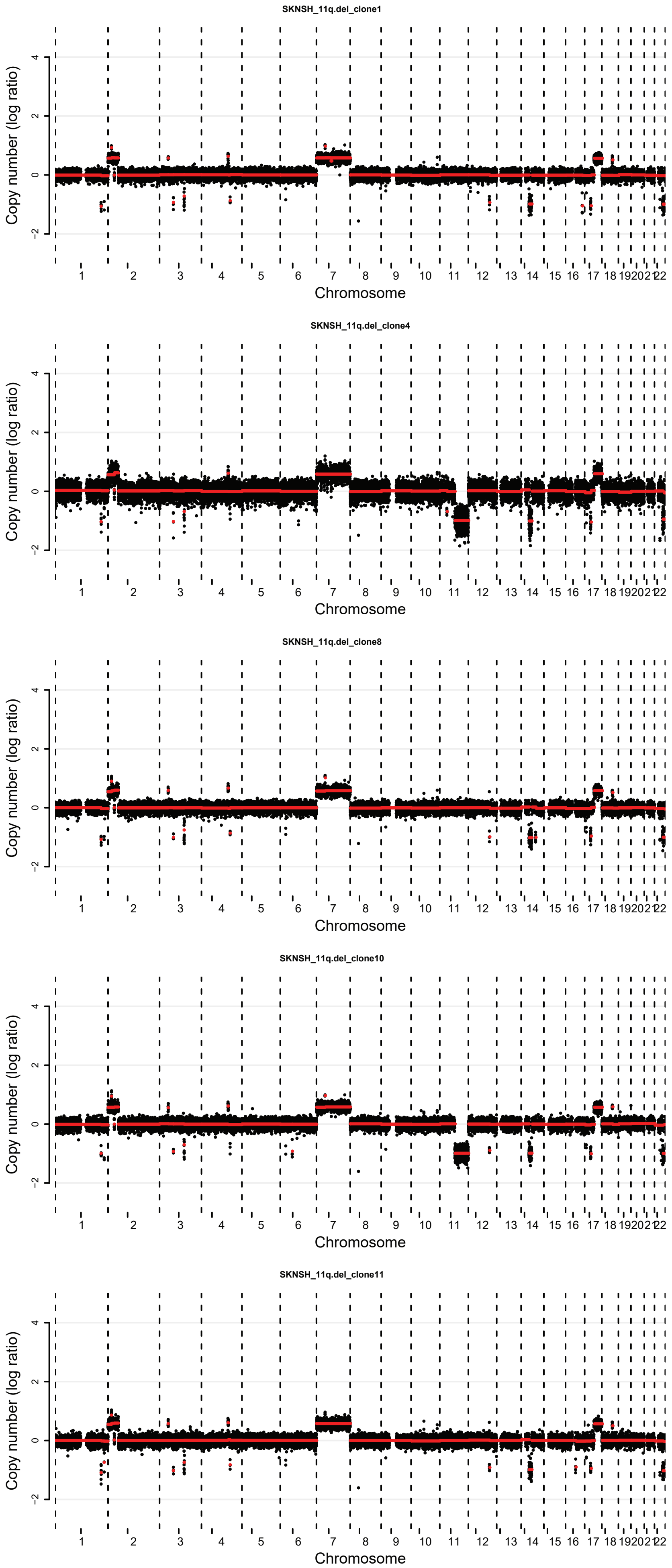


**Supplementary Figure 4:** Complementary to Figure 3A. Copy number profiles of translocation positive (clone 4, 10) and negative (clone 1,8,11) SKNSH clones. The x-axis shows the genomic location and the y-axis shows median-normalized log2-transformed copy number, with black dots representing bins and red lines representing segmented copy numbers.

### Supplementary Figure 5

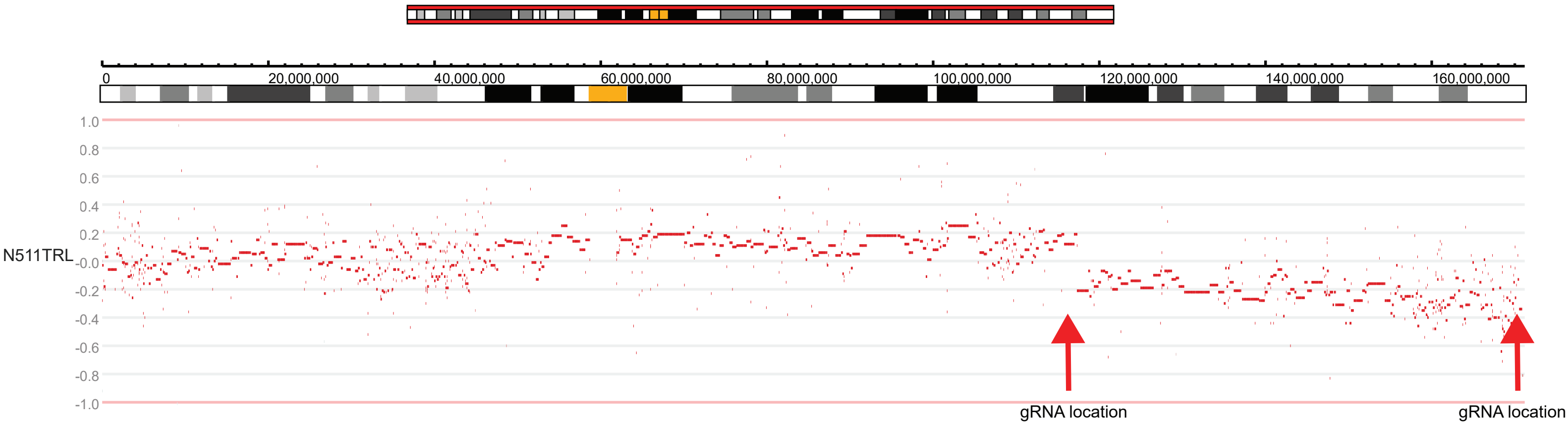


**Supplementary Figure 5:** Copy number profile of neuroblastoma tumor with a 6q deletion. The red arrows indicates the position of the 6q MID and 6q TERM gRNAs. Data was generated by WGS (PMID:26121087) and visualized using the R2 bioinformatics platform. Red dots represent segmented values. Ideograms representing the chromosomal location and banding pattern (centromeres are represented in yellow) are shown above the profile.

### Supplementary Figure 6

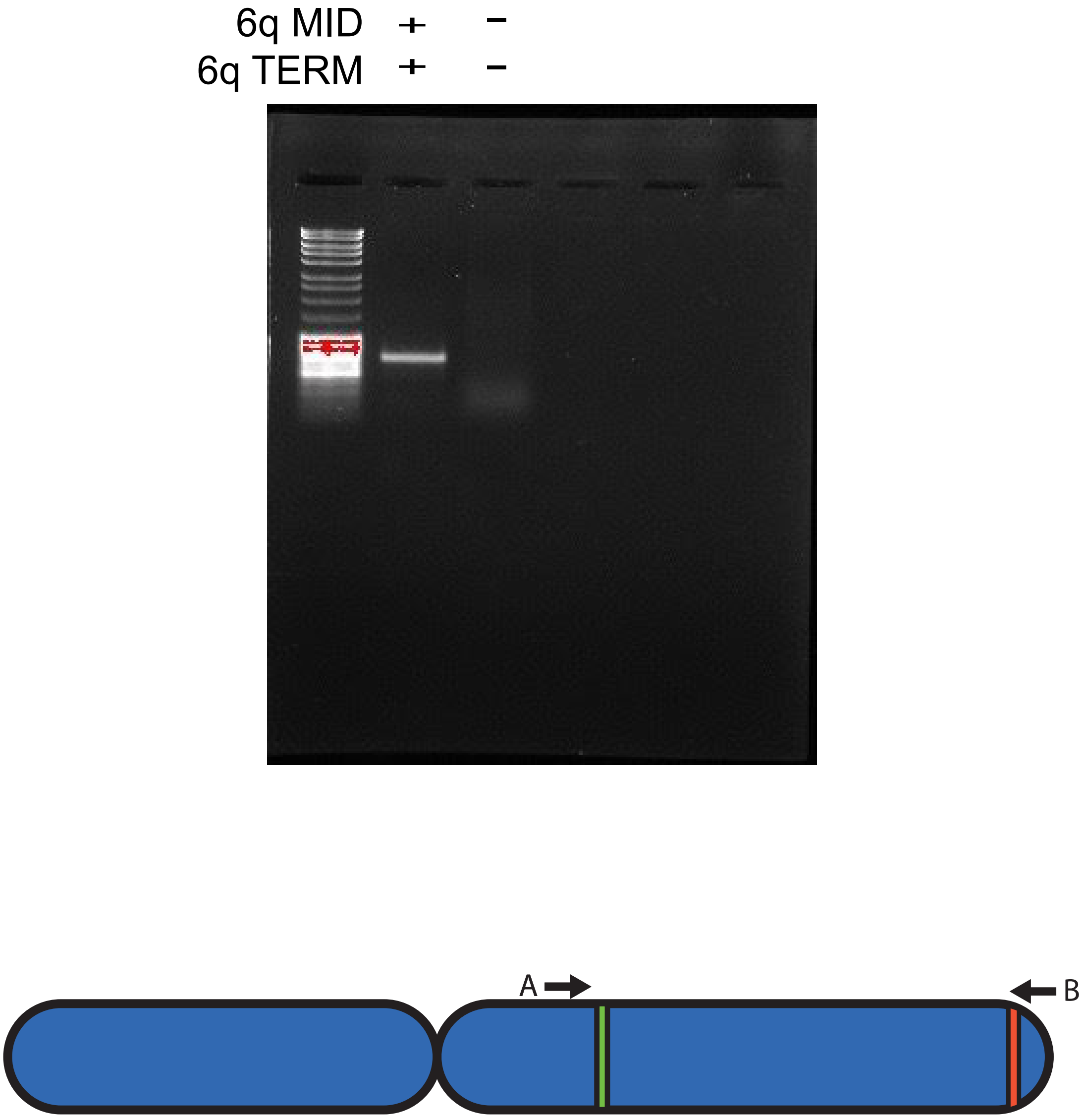


**Supplementary Figure 6:** Agarose gel electropohoresis of PCR reactions performed on cells transfected with 2 gRNAs  (6q MID and 6q TERM, lane 2), and wild type cells (lane 3). Lane 1 contains Generuler 1kb DNA ladder (Thermo Fisher Scientific). The primers used are shown in the lower figure.

### Supplementary Figure 7

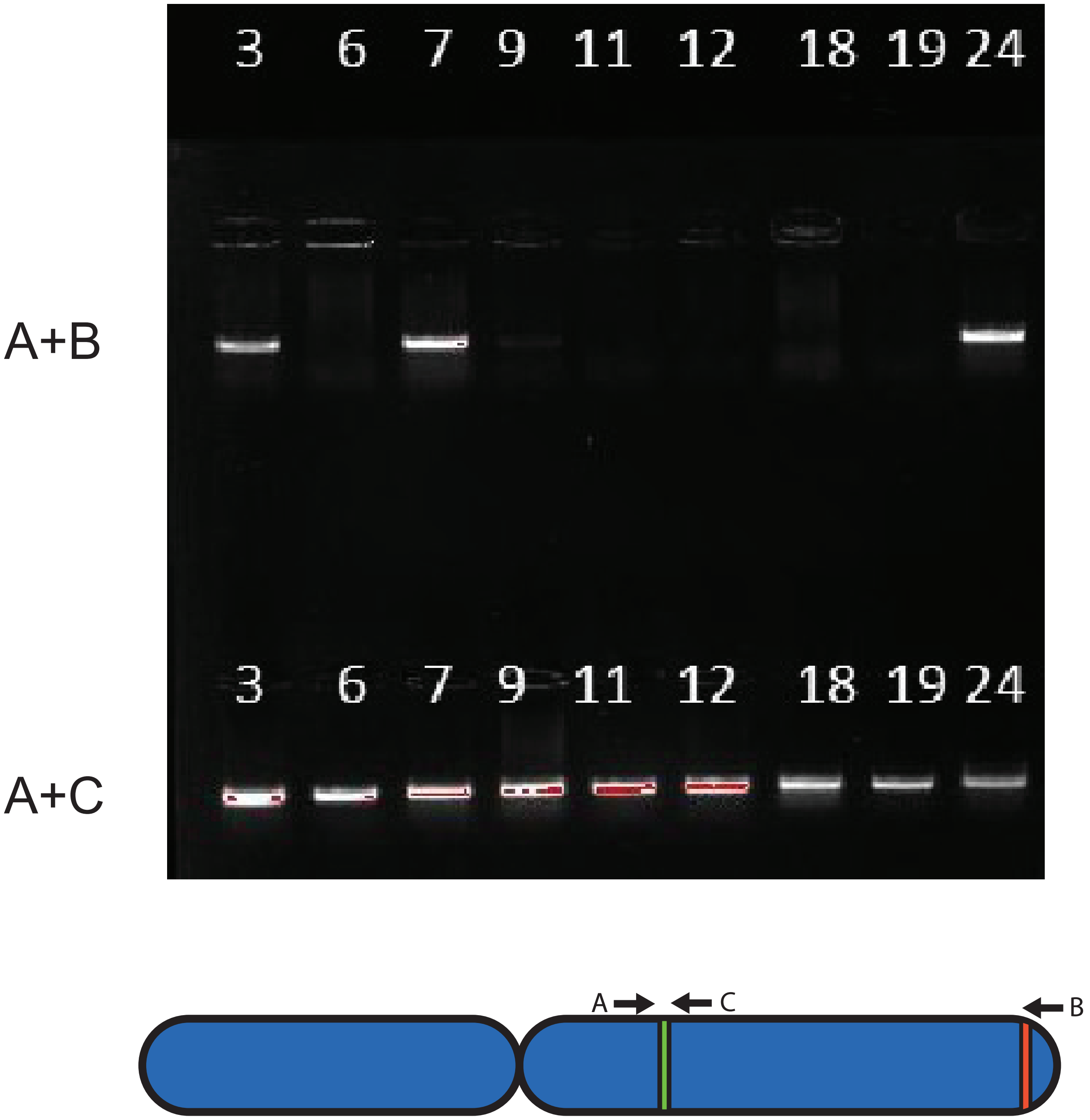


**Supplementary Figure 7:** Agarose gel electropohoresis of PCR reactions performed on clones derived from NMB cells transfected with 2 gRNAs for 6q. Primers used are indicated in front of the gel image, Lane 1 contains Generuler 1kb DNA ladder (Thermo Fisher Scientific) primer locations are shown in the lower figure.

### Supplementary Figure 8

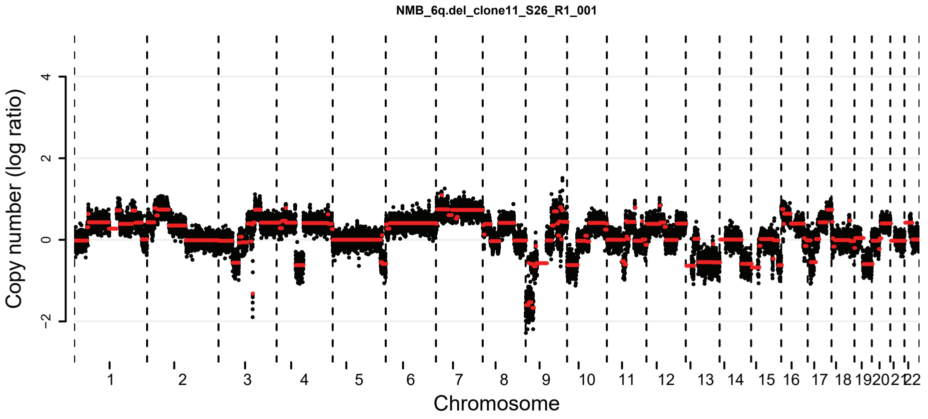


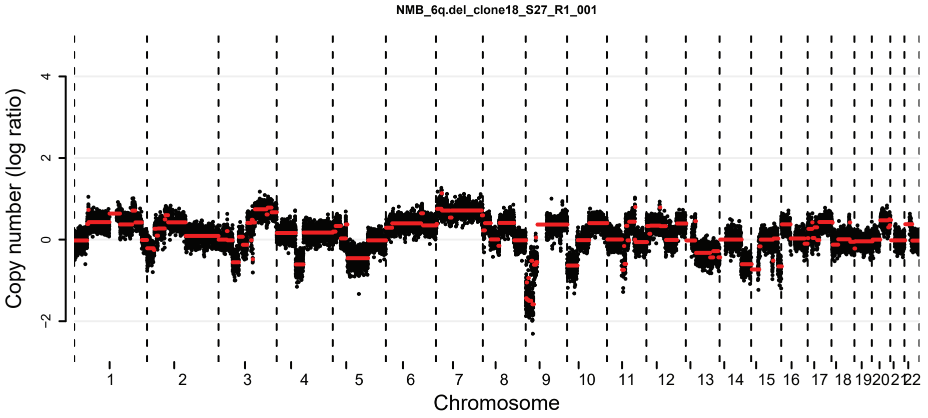


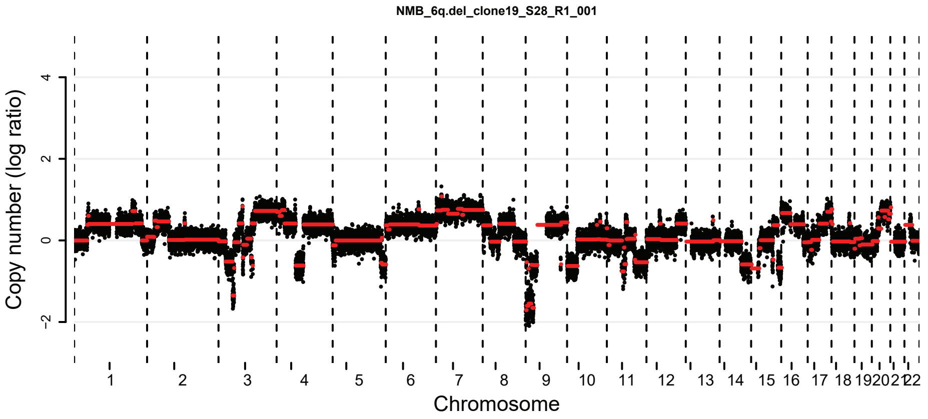


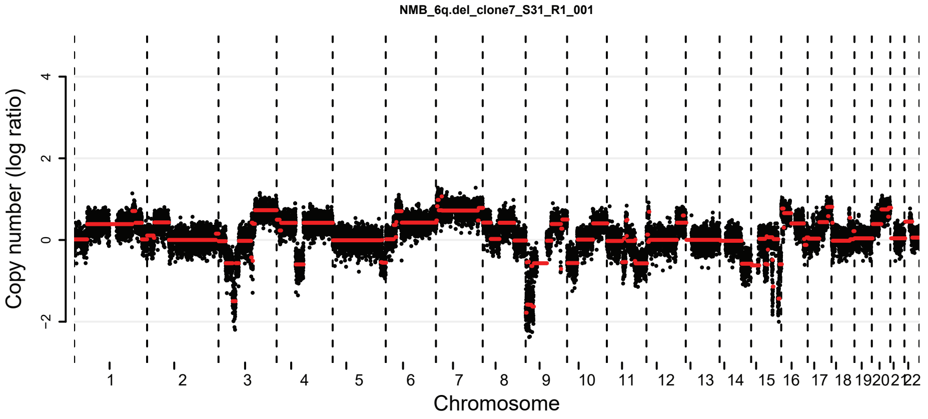


**Supplementary Figure 8:** Complementary to Figure 4A. Copy number profiles of translocation positive (clone 7) and negative (clone 11,18,19) NMB clones. The x-axis shows the genomic location and the y-axis shows median-normalized log2-transformed copy number, with black dots representing bins and red lines representing segmented copy numbers.
